# A Drd1-cre mouse line with nucleus accumbens gene dysregulation exhibits blunted fentanyl seeking

**DOI:** 10.1101/2025.02.14.638324

**Authors:** Annalisa Montemarano, Logan D. Fox, Farrah A. Alkhaleel, Alexandria E. Ostman, Hajra Sohail, Samiksha Pandey, Megan E. Fox

## Abstract

The synthetic opioid fentanyl remains abundant in the illicit drug supply, contributing to tens of thousands of overdose deaths every year. Despite this, the neurobiological effects of fentanyl use remain largely understudied. The nucleus accumbens (NAc) is a central locus promoting persistent drug use and relapse, largely dependent on activity of dopamine D1 receptors. NAc D1 receptor-expressing medium spiny neurons (D1-MSNs) undergo molecular and physiological adaptations that contribute to negative affect during fentanyl abstinence, but whether these neuroadaptations also promote fentanyl relapse is unclear. Here, we obtained Drd1-cre^120Mxu^ mice to investigate D1-dependent mechanisms of fentanyl relapse. We serendipitously discovered this mouse line is resistant to fentanyl seeking, despite similar intravenous fentanyl self-administration, and greater fentanyl-induced locomotion, compared to wildtype counterparts. In drug naïve mice, we found Drd1-cre^120Mxu^ mice have elevated D1 receptor expression in NAc, alongside increased expression of MSN marker genes *Chrm4* and *Penk*. We show Drd1-cre^120Mxu^ mice have increased sensitivity to the D1 receptor agonist SKF-38393, and exhibit divergent expression of MSN markers, opioid receptors, glutamate receptor subunits, and TrkB after fentanyl self-administration that may underly blunted fentanyl seeking. Finally, we show fentanyl-related behavior is unaltered by chemogenetic manipulation of D1-MSNs in Drd1-cre^120Mxu^ mice. Conversely, chemogenetic stimulation of putative D1-MSNs in wildtype mice recapitulated the blunted fentanyl seeking of Drd1-cre^120Mxu^ mice, supporting a role for aberrant D1-MSN signaling in this behavior. Together, our data uncover alterations in NAc gene expression and function with implications for susceptibility and resistance to developing fentanyl use disorder.

## Introduction

Opioid use disorder (OUD) is a chronic, relapsing condition characterized by uncontrollable opioid use and craving despite negative consequences, and the development of tolerance and withdrawal [1]. In 2022, over 9 million adults in the U.S. required OUD treatment [2], and over 80,000 individuals died from fatal overdose involving opioids [3]. A massive contributor to this epidemic is the current abundance of fentanyl in the illicit drug supply [4]. Fentanyl is a potent synthetic opioid that possesses unique pharmacological properties compared to other opioids: it has high lipid solubility that allows for rapid passage through the blood brain barrier [5], unique binding interactions with the mu opioid receptor [6], and higher selectivity for mu over kappa and delta opioid receptors as compared to morphine [7]. Fentanyl exposure is associated with poorer treatment outcomes for OUD [8], necessitating novel treatment approaches to cases of OUD involving fentanyl.

The vicious cycle of OUD, in which an abstinence-induced negative affective state promotes persistent relapse, is thought to be driven by neural circuitry centered around the nucleus accumbens (NAc) [1,9]. The NAc, located in the ventral striatum, is primarily comprised of GABAergic medium spiny neurons (MSNs) that are divided into two subtypes depending on whether they express *Drd1* (encoding dopamine D1 receptor; D1-MSNs) or *Drd2* (encoding dopamine D2 receptor; D2-MSNs). The MSN subtypes can be further identified by co-expression of additional markers: D1-MSNs co-express *Chrm4* (muscarinic acetylcholine receptor M4), *Pdyn* (preprodynorphin), and *Tac1* (preprotachkykinin-1); D2-MSNs co-express *Adora2a* (adenosine A_2A_ receptor), *Gpr6* (G protein-coupled receptor 6), and *Penk* (preproenkephalin) [10,11]. D1- and D2-MSNs have distinct projection targets [11–13] that typically drive opponent processes in reward behavior, where D1-MSN activity is classically considered “pro-reward” while D2-MSN activity is “anti-reward” [14–21], although there are numerous exceptions [22–29].

NAc MSNs undergo extensive physiological [30–42] and transcriptional [43–51] adaptations following opioid abstinence that are thought to promote negative affect and relapse. Like natural opioids, fentanyl self-administration produces a negative affective state [52,53] and an abstinence-induced withdrawal syndrome [54]. We recently showed NAc MSNs play unique roles during fentanyl abstinence: after homecage oral fentanyl, D1- but not D2-MSNs undergo transcriptional, morphological, and physiological adaptations that drive negative affect during abstinence [55]. However, whether these neuroadaptations generalize to intravenous fentanyl self-administration, or whether they promote fentanyl relapse, remains unknown.

To further understand how NAc D1-MSNs influence intravenous fentanyl self-administration and relapse to fentanyl seeking, we obtained a Drd1-cre mouse line deposited to the Jackson Laboratory [56] that has been used by several other groups [57–71]. First, we characterized baseline fentanyl self-administration and seeking behavior, and serendipitously discovered that the Drd1-cre^120Mxu^ (MMRC #037156-JAX) line demonstrates profound resistance to fentanyl seeking, despite similar acquisition and fentanyl intake as their wildtype littermates. Drd1-cre^120Mxu^ mice also show reduced fentanyl conditioned place preference, despite increased fentanyl-induced hyperlocomotion, as well as increased sensitivity to D1 receptor agonism. We further show these mice have differential mRNA expression of several NAc MSN markers both at baseline and after prolonged abstinence from fentanyl self-administration, including elevated *Drd1* expression at both timepoints. Lastly, we show that chemogenetic stimulation of putative NAc D1-MSNs in wildtype mice recapitulates the blunted fentanyl seeking exhibited by Drd1-cre^120Mxu^ mice, supporting aberrant signaling in NAc D1-MSNs as a mechanism promoting fentanyl resistance in the Drd1-cre^120Mxu^ line.

## Materials and Methods

### Experimental subjects

All procedures were approved by the Institutional Animal Care and Use Committee at the Pennsylvania State University College of Medicine (PSUCOM) and conducted in accordance with NIH guidelines for the use of laboratory animals. Drd1-cre^120Mxu^ mice on a C57BL/6J background were obtained from The Jackson Laboratory (B6;129-Tg(Drd1-cre)120Mxu/Mmjax; Jax strain #024860; RRID:MMRRC_037156-JAX) [56]. At PSUCOM, Drd1-cre^120Mxu^ hemizygotes were bred with C57BL/6J wildtype mice, also obtained from The Jackson Laboratory. Mice were given food and water *ad libitum* and housed in the PSUCOM vivarium on a 12:12h light:dark cycle with lights on at 7:00. All mice were housed in corncob bedding and provided with nestlets. Mice were housed in groups of 2-5, until they received intravenous surgery, after which they were pair-housed across a perforated acrylic divider throughout the entirety of self-administration.

### Procedures

All procedures are detailed in Supplemental.

### Statistics

Data were analyzed with Graphpad Prism 10 (La Jolla, CA) and JASP [72]. In the absence of significant sex effects or interactions, we collapsed data by sex. Any repeated measures data (e.g. self-administration, locomotion) were analyzed with repeated-measures ANOVA (RM-ANOVA) employing a Greenhouse-Geisser sphericity correction, with sex and genotype as between-subjects factors. Drug seeking data were analyzed with ANOVA using sex and genotype as between-subjects factors, and nose-poke as within-subject factor. Data without repeated measures (e.g. conditioned place preference, stereotypy) were analyzed by unpaired t-test due to no sex effects, or ANOVA when sex was significant. Gene expression data were analyzed with 2-way ANOVA using sex and genotype as between-subjects factors. For chemogenetic experiments, seeking was analyzed with ANOVA using sex, genotype, and virus as between factors, and nose-poke as within-subject factor. C-fos mRNA expression was analyzed with Welch’s ANOVA due to unequal variance across the groups. All post-hoc tests employed Sidak’s correction, except for the chemogenetic conditioned place preference and c-fos experiments which employed Dunnett’s correction comparing to the mCherry control.

## Results

NAc dopamine D1 receptors (D1R) play key roles in mediating reward and addiction phenotypes [9]. To answer questions about the mechanisms of D1-neurons in fentanyl use and relapse, we obtained Drd1-cre^120Mxu^ [73] mice from the Jackson Laboratory. While this Drd1-cre line has been used by other investigators in recent papers [57–71], known issues in other lines [74–77] necessitate characterizing behavior of all transgenic mice. Thus, we first asked if Drd1-cre^120Mxu^ mice would self-administer intravenous fentanyl. Similar to our previous work in wildtype C57/BL6J mice [78], Drd1-cre positive and negative mice underwent 10d of fentanyl self-administration training (timeline in **Figure 1A**). Both genotypes self-administered a similar number of fentanyl infusions (**Figure 1B**, sexes combined due to no sex effects or interactions. RM-ANOVA, Day: F_3.030,415.2_=3.40, p=0.018; Day x Genotype: F_9,1233_=1.06, p=0.39). Both genotypes learned to discriminate between active and inactive nose-pokes as measured by discrimination index (**Figure 1C**, active minus inactive nose-pokes, RM-ANOVA, Day: F_2.975, 407.6_=6.27, p=0.0004; Day x Genotype: F_9,1233_=0.67, p=0.72). Across 10d of self-administration, both genotypes had similar total fentanyl intake (wildtype female: 0.24±0.02, male: 0.19±0.02; Drd1-cre female: 0.21±0.02, male: 0.22±0.04 mg/kg).

**Figure 1.**
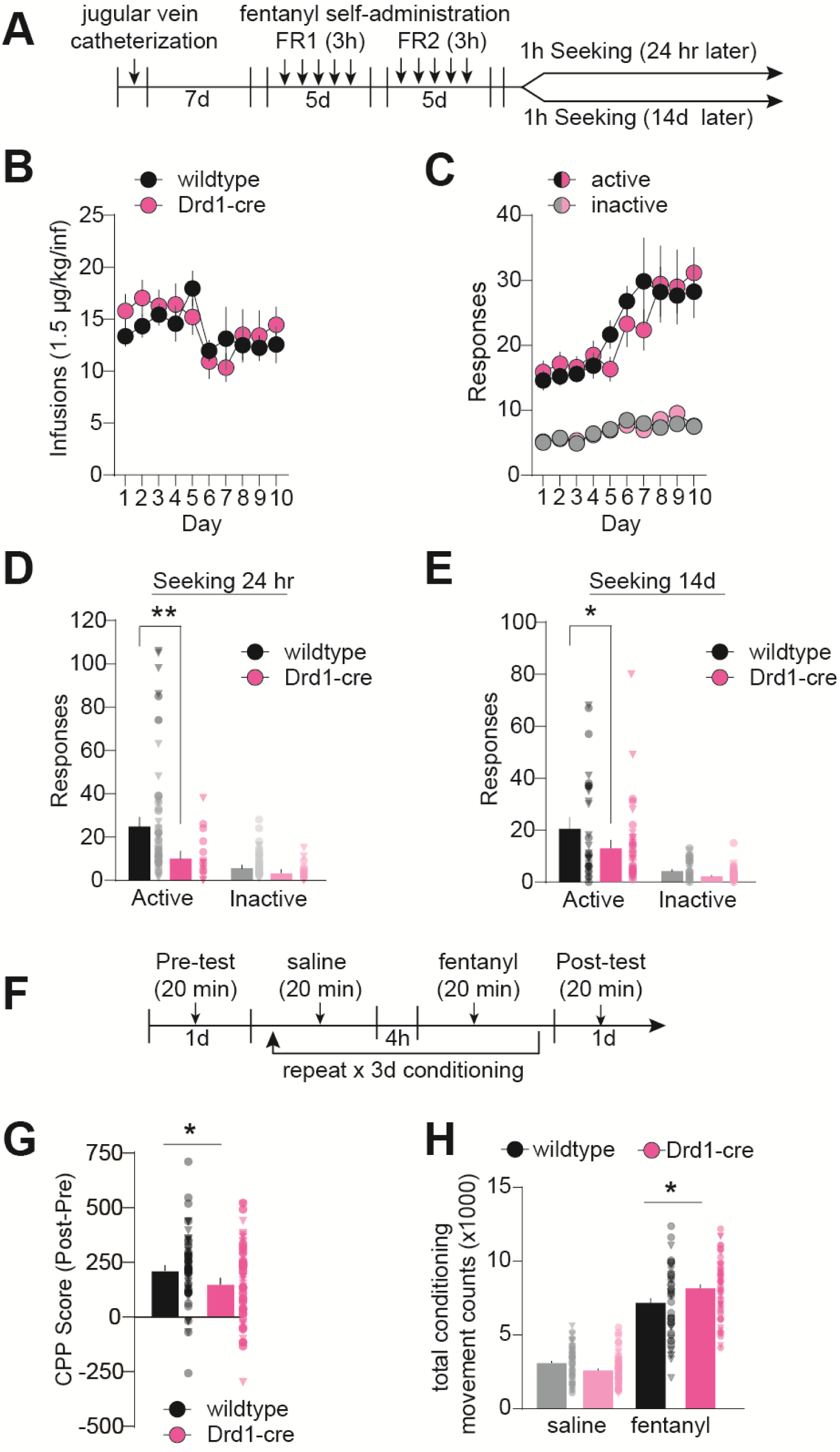
Drd1-cre^120Mxu^ mice show blunted fentanyl seeking relative to wildtype mice. (**A**) Experimental timeline for self-administration. Following recovery from jugular vein catheter surgery, mice underwent 5 days of fentanyl self-administration training (3 hr sessions) under fixed ratio 1 (FR1), followed by 5 days under FR2. Half of the mice underwent a non-reinforced seeking test 24hr after the last self-administration session and the other half underwent seeking 14d later. (**B**) Number of fentanyl infusions (1.5 µg/kg) earned during self-administration training in Drd1-cre^120Mxu^ (magenta) and wildtype (black) mice during training (wildtype n=38♀, 48♂; Drd1-cre^120Mxu^ n=27♀, 26♂). (**C**) Number of active and inactive responses during fentanyl self-administration training. (**D**) Number of active and inactive responses during a 1-hour non-reinforced seeking test in wildtype and Drd1-cre^120Mxu^ mice (**, p=0.026. Sidak’s post-hoc after 2-way ANOVA. Triangles denote datapoints from males and circles from females). (**E**) Number of active and inactive responses during a seeking test in separate mice after 14 days abstinence (*, p=0.03, Sidak’s post-hoc after 2-way ANOVA). (**F**) Experimental timeline for conditioned place preference. Mice freely explored the apparatus for 20 min during the pre-test day. For the following three days, mice received saline (10 ml/kg ip) in one compartment, followed 4h later by fentanyl (0.2 mg/kg ip) in the other. On the fifth day, mice freely explored the apparatus. (**G**) Preference for the fentanyl-paired compartment expressed as time spent in fentanyl-paired chamber during the post-test minus time spent during pre-test (*, p=0.03, unpaired t-test. Wildtype n=33♀, 30♂; Drd1- cre^120Mxu^ n=33♀, 32♂). (**H**) Total number of movement counts during conditioning sessions (*, p=0.0047, Sidak’s post-hoc after 2-way ANOVA). Data are presented as mean±SEM with individual mice overlaid.

We next asked if Drd1-cre^120MXu^ mice exhibited fentanyl seeking under extinction conditions. As expected, after 24hr abstinence, wildtype mice exhibited fentanyl seeking behavior marked by increased responding at the active nose-poke. By contrast, Drd1-cre^120Mxu^ mice exhibited virtually no fentanyl seeking behavior (**Figure 1D**, 2-way ANOVA, Genotype: F_1,71_=5.2, p=0.02; Active nose-poke, wildtype vs Drd1-cre, p=0.006, Sidak’s). We tested a separate set of mice after 14d abstinence. Like the 24hr timepoint, wildtype, but not Drd1-cre^120Mxu^ mice exhibited fentanyl seeking behavior (**Figure 1E**, 2-way ANOVA, Genotype: F_1,64_=4.23, p=0.04; Active nose-poke, wildtype vs Drd1-cre^120Mxu^, p=0.03, Sidak’s). To see if this generalized to other forms of fentanyl reward, we used conditioned place preference (CPP) in a separate set of mice (Timeline in **Figure 1F**). We found slightly reduced CPP scores in Drd1-cre^120Mxu^ mice relative to wildtype (**Figure 1G**, t_1,126_=2.17, p=0.03). Strikingly, when we examined fentanyl-induced hyperlocomotion, another proxy for drug reward, we found Drd1-Cre^120Mxu^ mice exhibit *greater* movement counts during fentanyl conditioning (**Figure 1H**, drug x genotype: F_1,109_=13.22 p=0.0004; wildtype vs Drd1-cre^120Mxu^ fentanyl, p=0.0047, Sidak’s). Finally, to determine if reduced drug reward was restricted to fentanyl, we asked if this generalized to other drugs with misuse potential, and compared cocaine self-administration and seeking in Drd1-cre^120Mxu^ mice relative to wildtype mice in our published dataset [79] (Timeline in **Supplemental Figure 1A**). Unlike fentanyl, Drd1-cre^120Mxu^ had reduced cocaine intake relative to wildtype mice (**Supplemental Figure 1B**, RM-ANOVA, Genotype: F_1,24_=10.8; **Supplemental Figure 1C,** Active responses: Genotype F_1,24_=17.3, p=0.0003). Like fentanyl, Drd1-cre^120Mxu^ had fewer active responses during the non-reinforced seeking test at 24h (**Supplemental Figure 1D,** Genotype: F_1,48_=22.6, p<0.0001; Active responses, wildtype vs Drd1-cre^120Mxu^, p=0.0001, Sidak’s).

Previous work using Drd2-EGFP mice found elevated striatal *Drd2* expression with effects on behavioral responses to cocaine [74]. Thus, we next asked if Drd1-cre^120Mxu^ mice also have altered baseline expression of dopamine receptors that might drive the behavioral phenotypes. Given the importance of NAc in opioid-reward and opioid seeking [1], we looked at gene expression in NAc. In experimentally naïve mice, we found elevated *Drd1* expression in both sexes of Drd1-cre^120Mxu^ mice compared to wildtype (females: 1.59±0.49, males: 1.60±0.14; **Figure 2A**, 2-way ANOVA, Genotype: F_1,19_=6.46, p=0.02). We also found elevated expression of D1- MSN marker *Chrm4* (**Figure 2B**, Genotype: F_1,19_=6.16, p=0.02). Not all D1-MSN markers were elevated, as there were no differences in *Pdyn* (**Figure 2C**), and only baseline sex differences in *Tac1* (**Figure 2D**, Sex: F_1,19_=6.1, p=0.023). We found no changes in D2-MSN markers *Drd2*, *Adora2a,* or *Gpr6* (**Figure 2E-G**), but a significant increase in *Penk* in Drd1-cre^120Mxu^ mice (**Figure 2H**, Genotype: F_1,19_=4.6, p=0.046). These differences in MSN marker genes did not generalize to all Drd1-cre mice, as we found no aberrant expression in a different Drd1- cre line (GensatFK150; **Supplemental Figure 1E**). Given the differences in fentanyl seeking between genotypes, we also looked at expression of opioid receptors, and the related nociceptin receptor in NAc (**Figure 2I-L**). In experimentally naïve mice, we found no differences in opioid receptor expression between genotypes, and only baseline sex differences in *Oprk1* expression (**Figure 2K**, Sex: F_1,19_=4.9, p=0.04).

**Figure 2.**
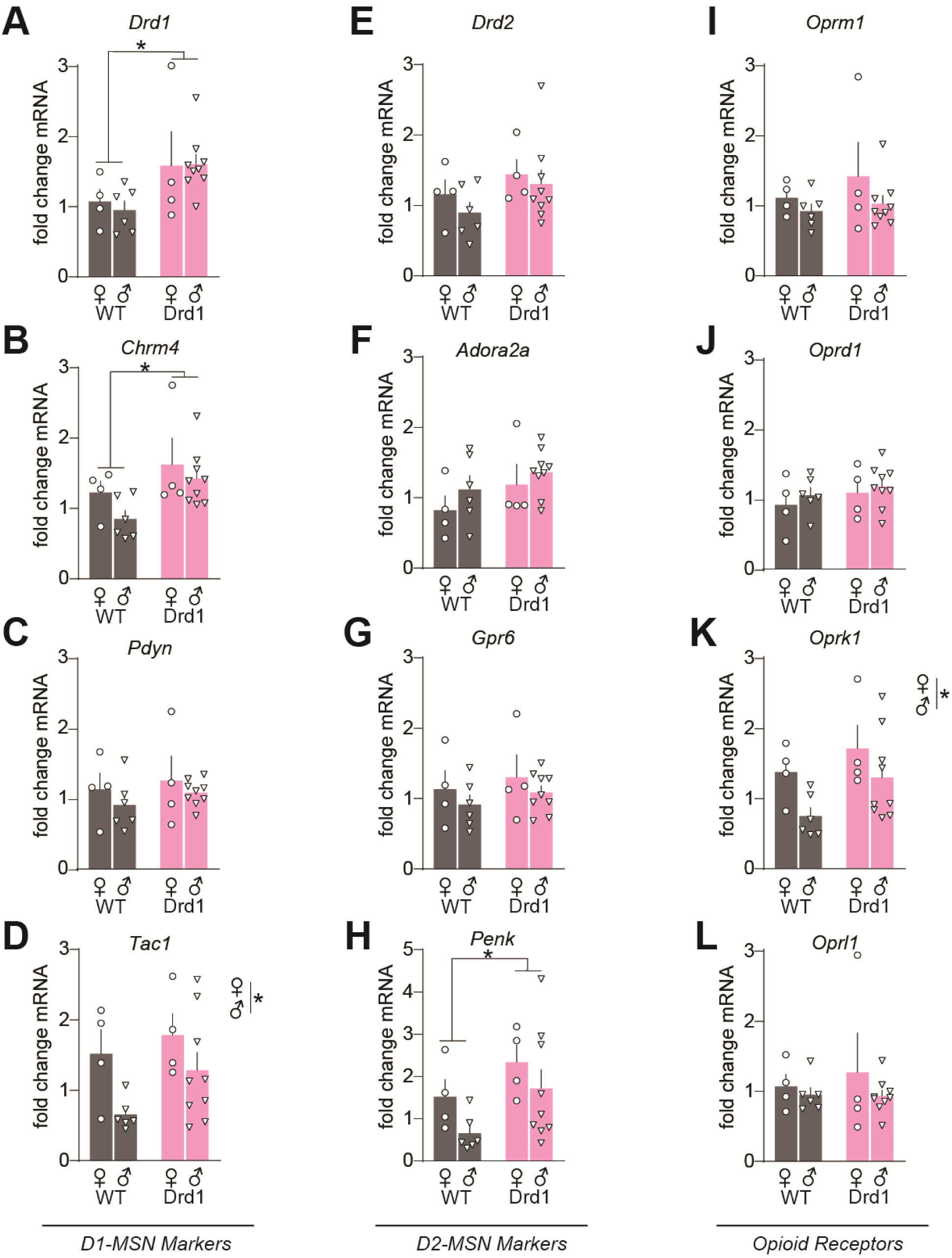
Drd1-cre^120Mxu^ mice have elevated expression of medium spiny neuron marker genes in nucleus accumbens at baseline. Fold change mRNA in experimentally naïve wildtype (black) and Drd1-cre^120Mxu^ (magenta) mice, relative to average of naïve male and female wildtype mice (wildtype n=4♀, 6♂; Drd1-cre^120Mxu^ n=4♀, 9♂). Data are presented as mean±SEM. (**A**) Dopamine D1 receptor, *, p=0.019, main effect of genotype. (**B**) Muscarinic acetylcholine receptor M4, *, p=0.022, main effect of genotype. (**C**) Preprodynorphin. (**D**) Preprotachykinin-1, *, p=0.023, main effect of sex. (**E**) Dopamine D2 receptor. (**F**) Adenosine A2a receptor. (**G**) G protein-coupled receptor 6. (**H**) Preproenkephalin, *, p=0.045, main effect of genotype. (**I**) Mu opioid receptor. (**J**) Delta opioid receptor. (**K**) Kappa opioid receptor, *, p=0.040, main effect of sex. (**L**) Opioid related nociceptin receptor 1. Data are mean±SEM with individual mice overlaid.

Since Drd1-cre^120Mxu^ mice had elevated *Drd1* expression in NAc, we next asked if this would produce a baseline locomotor phenotype or increased sensitivity to D1R agonists. We administered D1R agonist SKF- 38393 (SKF, 30 mg/kg sc), or saline, just prior to open field testing, which typically produces hyperlocomotion in unhabituated mice[80]. Both SKF-treated wildtype and Drd1-cre^120Mxu^ mice exhibited increased locomotion relative to their saline-treated counterparts (**Figure 3A**, time x drug: F_4.9,186.4=_7.39, p<0.001). Contrary to our prediction of increased distance traveled, both saline- and SKF-treated Drd1-cre^120Mxu^ traveled less total distance across the 90 minutes relative to wildtype (Genotype: F_1,38_=4.1, p=0.049), that was not mediated by differences in average velocity (**Figure 3B**). Given the baseline differences in locomotion, we next looked at distance traveled as a percent of saline treatment. Drd1-cre^120Mxu^ mice showed greater SKF-potentiated locomotion relative to wildtype mice, suggesting increased D1R sensitivity (**Figure 3C**, t_1,21_=2.79, p= 0.011). Since D1R activation can elicit stereotyped movements [81], we also looked at jumps and stationary movement counts (stereotypy counts). Drd1-cre^120Mxu^ mice exhibited more SKF-induced jumps (**Figure 3D**, t_1,21_=3.33, p=0.003) and stereotypy counts (**Figure 3E**, t_1,21_=5.9 p<0.0001). Together, these data suggest Drd1-cre^120Mxu^ mice have reduced locomotion in a novel environment, but increased sensitivity to D1R activation.

**Figure 3.**
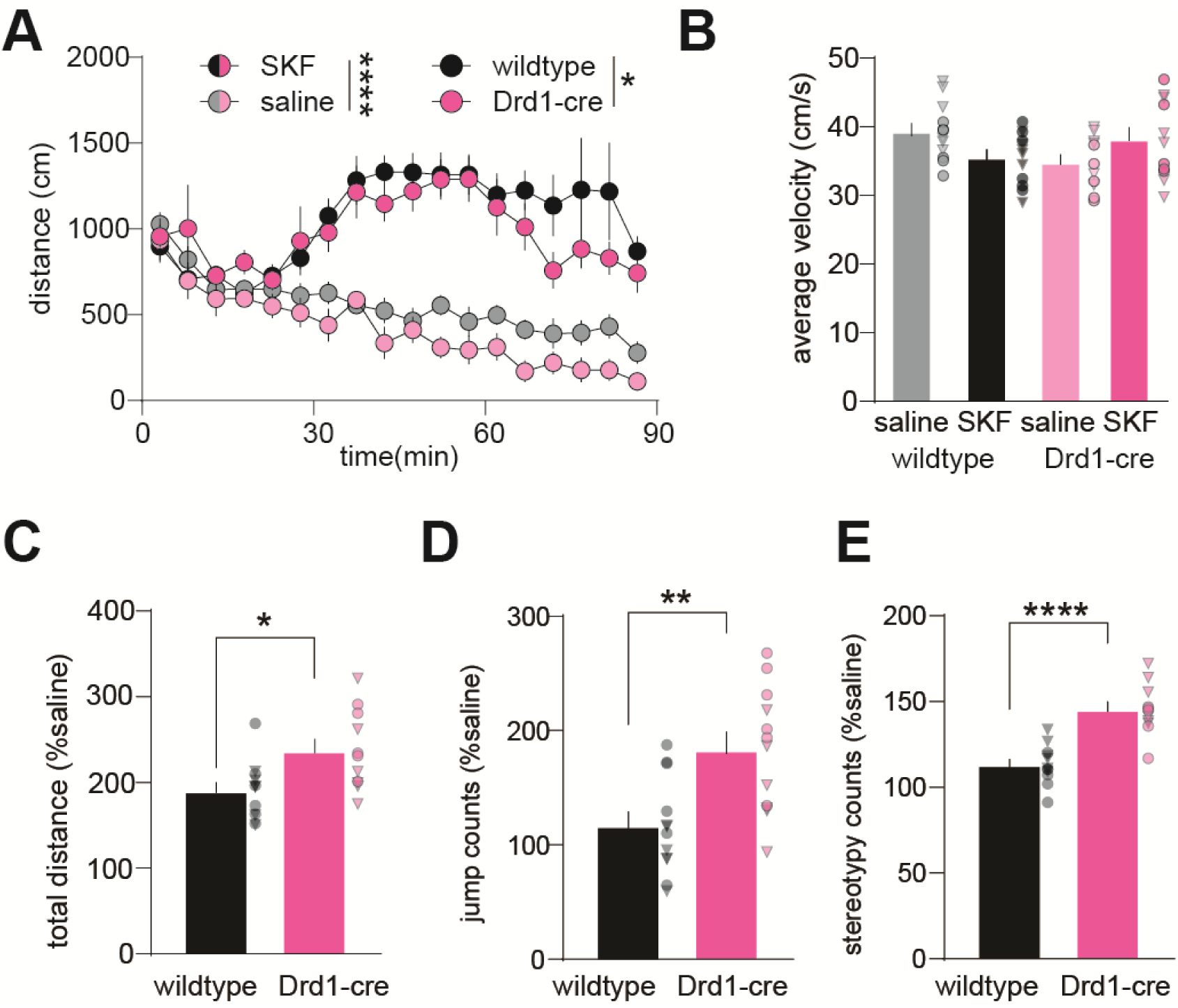
Drd1-cre^120Mxu^ mice have attenuated locomotor response to a novel environment and show increased sensitivity to a D1 receptor agonist. Male and female wildtype (black) and Drd1-cre^120Mxu^ (magenta) mice received saline or 30 mg/kg SKF-38393 s.c. prior to open field testing (wildtype saline n=6♀, 6♂; wildtype SKF n=6♀, 6♂; Drd1-cre saline n=5♀, 6♂; Drd1-cre SKF n=5♀, 6♂). (**A**) Distance traveled in the open field over 90 min in 5min bins. ****, p<0.0001, main effect of drug; *, p=0.044, main effect of genotype. (**B**) Average velocity in the open field in saline and SKF treated mice. Triangles represent data points from male mice and circles from females. (**C**) Total distance traveled in SKF-treated wildtype and Drd1-cre relative to saline-treated mice. *, p= 0.011, unpaired t-test. (**D**) Jump counts in SKF-treated mice relative to saline-treated mice. **, p=0.003, unpaired t-test. (**E**) Stereotypy counts in SKF-treated mice relative to saline-treated mice. ****, p<0.0001, unpaired t-test. Data are presented as mean±SEM with individual mice overlaid.

Given the increase in *Drd1* expression, and increased response to a D1R agonist in experimentally naïve Drd1-cre^120Mxu^ mice, we next asked if elevated *Drd1* expression was maintained in mice with fentanyl self-administration experience, as this may underlie their behavioral differences (Timeline in **Figure 4A**). After 14d abstinence, fentanyl-experienced Drd1-cre^120Mxu^ mice maintained elevated *Drd1* expression relative to naïve wildtype mice (Drd1-cre females: 1.25±0.09, males: 1.7±0.16; **Figure 4B**, Sex x Genotype: F_1,17_=8.78, p=0.009; females: p=0.0005, males: p<0.0001, Sidak’s). This is juxtaposed against a relative *Drd1* downregulation in fentanyl-experienced wildtype mice (wildtype females: 0.69 ±0.08, males: 0.62 ± 0.04; **Figure 4B**). We also looked beyond *Drd1*, and a drastic genotype difference emerged that was not present in naïve mice.

**Figure 4.**
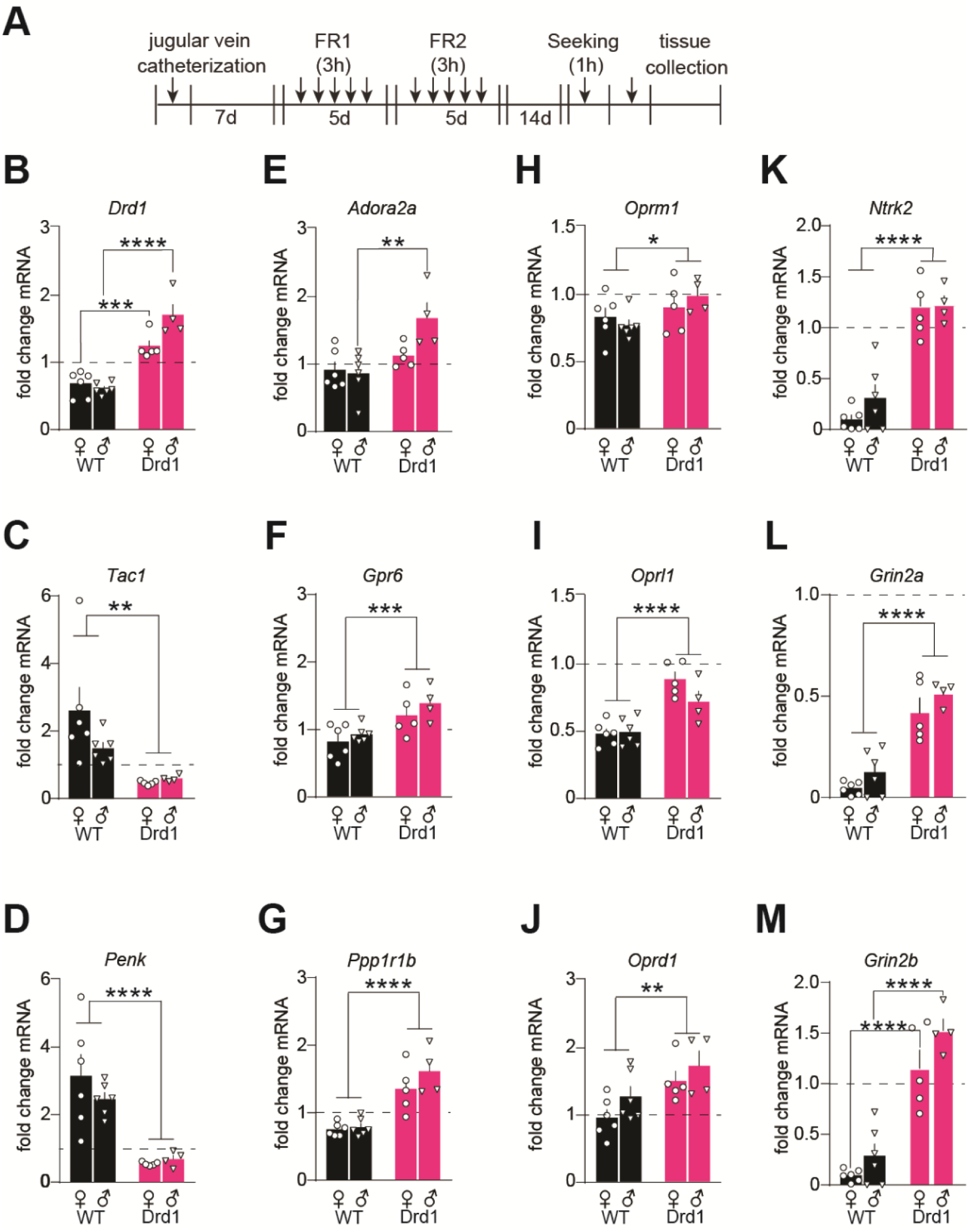
Molecular adaptations in nucleus accumbens differ across genotype following abstinence from fentanyl self-administration. (**A**) Experimental timeline. Following recovery from jugular vein catheter surgery, mice underwent 5 days of fentanyl self-administration under fixed ratio 1 (FR1), followed by 5 days under FR2. Following 14 days of homecage abstinence, mice underwent a non-reinforced seeking test, then nucleus accumbens tissue was collected. (**B-M**) Fold change mRNA relative to experimentally naïve male and female wildtype mice (dashed line; wildtype n=6♀, 6♂; Drd1-cre^120Mxu^ n=5♀, 4♂). Data are presented as mean±SEM. (**B**) Dopamine D1 receptor, Sex x Drug Interaction, p=0.0087; ***p=0.0005, ****p<0.0001, wildtype vs Drd1-cre females, males, respectively, Sidak’s post-hoc. (**C**) Preprotachykinin-1, **p=0.0021, main effect of genotype. (**D**) Preproenkephalin, ****p<0.0001, main effect of genotype. (**E**) Adenosine A2a, Sex x Drug Interaction, p=0.044; **p=0.0018, wildtype vs Drd1-cre males, Sidak’s post-hoc. (**F**) G protein-coupled receptor 6, ***p=0.001, main effect of genotype. (**G**) Dopamine and cAMP- regulated phosphoprotein (DARPP-32), ****p<0.0001, main effect of genotype. (**H**) Mu opioid receptor, *p=0.041, main effect of genotype. (**I**) Opioid related nociceptin receptor 1, ****p<0.0001, main effect of genotype. (**J**) Delta opioid receptor, **p=0.005, main effect of genotype. (**K**) Tropomyosin receptor kinase B (TrkB), **** p<0.0001, main effect of genotype. (**L**) NMDA receptor Subunit 2A, ****p<0.0001, main effect of genotype. (**M**) NMDAR Subunit 2B, ****p<0.0001, wildtype vs Drd1-cre females, males, respectively, Sidak’s post-hoc. Data are presented as mean±SEM with individual mice overlaid.

Wildtype females had even greater *Tac1* expression at the 14d abstinence timepoint compared to baseline (females: 2.6±0.069, males: 1.49 ±0.18; **Figure 4C**). By contrast, *Tac1* was downregulated in Drd1-cre^120Mxu^ mice with fentanyl experience (females: 0.47±0.029, males: 0.6±0.051; **Figure 4C**, Genotype: F_1,17_=13.10, p=0.002). In some cases, fentanyl experience normalized previous differences in gene expression between genotype (e.g. *Chrm4*, **Supplemental Figure 2C**). In other cases, differences in gene expression were reversed, such as reduced *Penk* expression in fentanyl-experienced Drd1-cre^120Mxu^ mice, (females: 0.54±0.02, males: 0.68±0.12; **Figure 4D**), and a relative upregulation in fentanyl-experienced wildtype mice (female: 3.15±0.06; male: 2.46±0.20; **Figure 4D**, Genotype: F_1,17_=30.7, p<0.0001). Surprisingly, D2-MSN marker genes *Adora2a*, and *Gpr6* were now upregulated in fentanyl-experienced Drd1-cre^120Mxu^ mice (**Figure 4E-F**, *Adora2a,* Sex x Genotype: F_1,17_=4.7, p=0.044, male wildtype vs Drd1-cre, p=0.0018; *Gpr6*, Genotype: F_1,17_=15.8, p=0.001), although *Adora2a* was only significantly upregulated in Drd1-cre^120Mxu^ males. We also found upregulation of general MSN marker *Ppp1r1b* (DARPP-32) in fentanyl-experienced Drd1-cre^120Mxu^ mice (**Figure 4G**, Genotype F_1,17_=43.5, p<0.0001), a difference that was absent in experimentally naïve mice (**Supplemental Figure 3A**). When we looked at opioid receptor expression, we again found differences that were absent in experimentally naïve mice. Compared to fentanyl-experienced wildtype mice, Drd1-cre^120Mxu^ mice had less downregulation of *Oprm1* (wildtype females: 0.82±0.07, males: 0.76±0.04; Drd1-cre females: 0.90±0.09, males 0.98±0.06; **Figure 4H**, Genotype: F_1,17_=4.9, p=0.04) and *Oprl1* (wildtype females: 0.48±0.04, males: 0.49±0.04; Drd1-cre females: 0.88±0.06, males: 0.71±0.08; **Figure 4I**, Genotype: F_1,17_=36.7, p<0.0001). Drd1-cre^120Mxu^, but not wildtype mice, also upregulated *Oprd1* (wildtype females: 0.95±0.12, males: 1.26±0.15; Drd1-cre females: 1.49±0.15, males: 1.72±0.22; **Figure 4J**, Genotype F_1,17_=10.2, p=0.0005). Because chronic opioid exposure is associated with reduction in NAc TrkB expression [17] and NAc NMDAR function [82], we also looked at expression of *Ntrk2*, and GluN2 subunits *Grin2a* and *Grin2b.* In fentanyl-experienced wildtype mice, we found expected downregulation of *Ntrk2* that was completely absent in Drd1-cre^120Mxu^ mice (wildtype female: 0.10±0.044, male: 0.31±0.013; Drd1-cre female: 1.17±0.12, male: 1.23±0.09; **Figure 4K**, Genotype: F_1,17_=88.25, p<0.0001). Similarly, downregulation of GluN2 subunits was more extensive in fentanyl-experienced wildtype mice compared with Drd1-cre^120Mxu^, especially for *Grin2b* (*Grin2a*: wildtype female 0.05±0.01, male 0.12±0.05; Drd1- cre female 0.43±0.07, male 0.51±0.03, Genotype F_1,17_=76.1, p<0.0001; *Grin2b*: wildtype female 0.09±0.02, male 0.29±0.012; Drd1-cre female 1.15±0.18, male 1.5±0.11, Genotype F_1,17_=91.0, p<0.0001; **Figure 4L-M**) None of these differences were present in experimentally naïve mice (**Supplemental Figure 3B-D**).

Due to persistent upregulation of NAc *Drd1* in the Drd1-cre^120Mxu^ mice, we next asked if chemogenetically manipulating activity of NAc D1-MSNs using Designer Receptors Exclusively Activated by Designer Drugs (DREADDs) could restore fentanyl seeking to that of wildtype mice (Timeline in **Figure 5A**). Immediately following fentanyl self-administration (**Figure 5B**), we assigned Drd1-cre^120Mxu^ mice to groups to ensure similar intake across sex and virus (future virus F_2,33_=0.4, future virus x sex F_1,33_=0.3). After two weeks of abstinence, we inhibited and activated NAc D1-MSNs with the DREADDs ligand deschloroclozapine (DCZ) prior to a fentanyl seeking test. Surprisingly, we found neither chemogenetic inhibition, nor activation, influenced fentanyl-seeking behavior in Drd1-cre^120Mxu^ mice (**Figure 5C**, nose-poke x virus F_2,35_=2.15, p=0.15).

**Figure 5.**
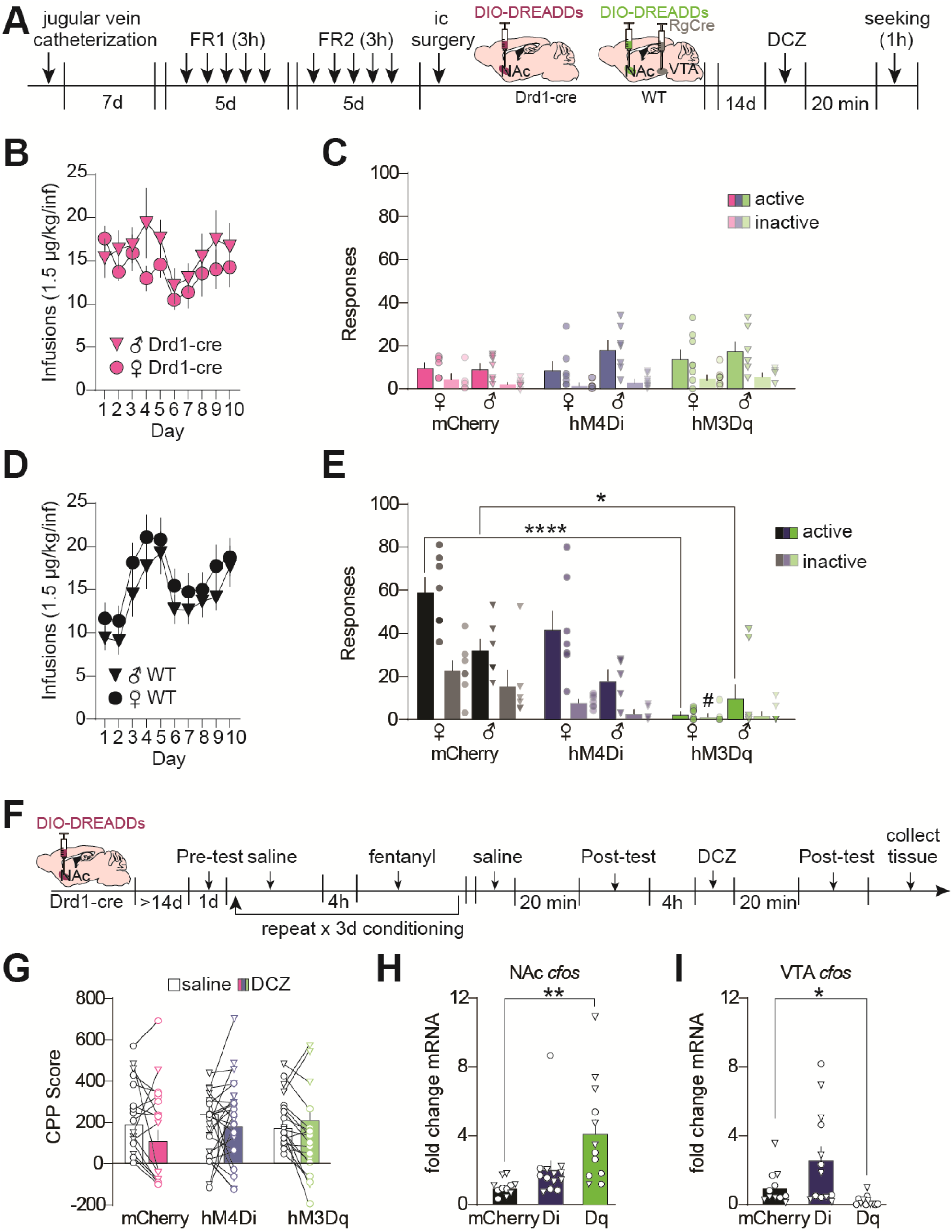
Chemogenetic manipulation of NAc D1-MSNs blunts fentanyl seeking in wildtype mice but does not alter seeking in Drd1-cre^120Mxu^ mice. (**A**) Experimental timeline for IVSA experiments. Following jugular vein catheter surgery mice underwent 10 days of fentanyl self-administration training. Mice then underwent surgery to express DREADDs in NAc D1-MSNs, using Cre-dependent DREADDs in NAc of Drd1-cre^120Mxu^ mice, or retrograde Cre in VTA and Cre-dependent DREADDs in NAc of wildtype mice. Following 14d abstinence and viral expression, mice were given 0.1mg/kg DCZ IP 20 min prior to a 1-hr seeking test. (**B**) Infusions earned during fentanyl self-administration in male and female Drd1-cre^120Mxu^ mice. (**C**) Active and inactive responses during the fentanyl seeking test in Drd1-cre^120Mxu^ mice expressing mCherry control (n=5♀, 7♂), inhibitory hM4Di (n=7♀, 7♂), or stimulatory hM3Dq (n=8♀, 6♂) in NAc D1-MSNs. (**D**) Fentanyl infusions earned during self-administration in male and female wildtype mice. (**E**) Active and inactive responses during the fentanyl seeking test in wildtype mice (mCherry, n=7♀, 6♂; hM4Di, n=7♀, 5♂; hM3Dq, n=7♀, 8♂). Active responses: ****p<0.0001, hM3Dq vs mCherry females; *p=0.024, hM3Dq vs mCherry males, Sidak’s. Inactive responses: #p=0.024, female hM3Dq vs mCherry. (**F**) Experimental timeline for CPP experiments. Drd1-cre^120Mxu^ mice underwent surgery for Cre-dependent DREADDs in NAc. Mice freely explored the apparatus during the pre-test day. For the following three days, mice received saline (10 ml/kg ip) in one compartment, followed 4h later by fentanyl (0.2 mg/kg ip) in the other. On the fifth day, mice received saline (or DCZ) 20 minutes prior to the first post-test. Then, 4h later, mice received 0.1 mg/kg DCZ (or saline) 20 minutes prior to the second post-test. Tissue was collected immediately following the second post-test. (**G**) CPP score during the post-test under saline and DCZ conditions (mCherry, n=12♀, 13♂; hM4Di, n=12♀, 7♂; hM3Dq, n=10♀, 6♂). (**H**) In mice receiving DCZ just prior to tissue collection, immediate early gene cfos is upregulated in NAc of Drd1-cre^120Mxu^ mice expressing hM3Dq relative to mCherry, **p=0.007, Dunnet’s T3 (mCherry, n=5♀, 4♂; hM4Di, n=7♀, 6♂; hM3Dq, n=8♀, 4♂). (**I**) In downstream VTA, cfos is downregulated in hM3Dq relative to mCherry, *p=0.049, Dunnet’s T3. Data are presented as mean±SEM with individual mice overlaid.

By contrast, chemogenetic activation of putative D1-MSNs suppressed fentanyl-seeking in wildtype mice (Self-administration in **Figure 5D**, seeking in **Figure 5E**; Nose-poke x sex x virus F_2,34_=3.44, p=0.044; Active nose-pokes, hM3Dq vs mCherry, male p=0.024, female p<0.0001, Sidak’s). Importantly, this was not due to differences in self-administration behavior, as all mice, regardless of genotype or sex, had comparable fentanyl intake during acquisition (wildtype female 0.26±0.02, male 0.21±0.02, Drd1-cre female 0.21±0.02, male 0.27±0.04 mg/kg, Genotype F_1,76_=0.03). As with fentanyl-seeking after self-administration, we found Drd1- cre^120Mxu^ mice were also insensitive to chemogenetic manipulation during fentanyl conditioned-place preference. Using a within-subject design (Timeline in **Figure 5F**), we found neither inhibiting, nor activating D1-MSNs significantly altered time spent in the fentanyl-paired chamber (**Figure 5G**, drug x virus F_2,54_=2.8, p=0.07). Importantly, this was not due to DCZ inefficacy in Drd1-cre^120Mxu^ mice, as stimulating NAc D1-neurons increased *cfos* in NAc (**Figure 5H**, Welch’s ANOVA, F_2,16.51_=7.8 p=0.0042, mCherry vs hM3Dq p=0.007, Dunnet’s T3), and decreased *cfos* in downstream VTA as expected (**Figure 5I**, Welch’s ANOVA, F_2,15.86_=7.7, mCherry vs hM3Dq=0.049, Dunnet’s T3).

## Discussion

Here we report that the widely-used Drd1-cre^120Mxu^ mouse line [73] exhibits several behavioral and transcriptional differences compared to wildtype mice. We serendipitously discovered that this line is resistant to fentanyl seeking, as demonstrated by dramatically reduced responding in a nonreinforced seeking test following either 24 hours or 14 days of abstinence, despite normal fentanyl acquisition. This resistance translates to other fentanyl administration paradigms, as Drd1-cre^120Mxu^ mice also have reduced preference for a fentanyl-paired context, despite increased fentanyl-induced hyperlocomotion. These mice also exhibit baseline hypolocomotion in a novel environment, as well as relative increased behavioral response to D1R agonism. Although the mechanism underlying fentanyl resistance in these mice is not fully understood, key differences in gene expression in the Drd1-cre^120Mxu^ mice, alongside evidence that stimulating putative D1-MSNs in wildtype mice produces similar resistance to fentanyl seeking, implicates that this phenotype is, at least in part, due to aberrant signaling in D1-MSNs.

Here we showed that Drd1-cre^120Mxu^ mice have normal acquisition of intravenous fentanyl self-administration, but reduced cocaine self-administration. Our study is not the first to report on a transgenic line with abnormal drug self-administration behavior; for example, ChAT-Cre mice (B6.FVB(Cg)-Tg(Chat-cre)GM60Gsat/Mmucd; RRID:MMRRC_030869-UCD) are reported to have reduced self-administration of intravenous nicotine [75]. However, our findings are unique in that despite normal acquisition of fentanyl self-administration, Drd1-cre^120Mxu^ mice exhibited attenuated nonreinforced fentanyl seeking. This effect is strikingly robust: the blunted fentanyl seeking is observed at both 24 hours and 14 days after self-administration, it is also reflected in a reduced preference for the fentanyl-paired context in CPP, and the behavior persists in both paradigms despite bidirectional chemogenetic manipulation of D1-MSNs. Interestingly, we found that stimulating putative D1-MSNs in wildtype mice by chemogenetically targeting ventral tegmental area (VTA)-projecting NAc neurons attenuated fentanyl seeking to levels seen in the Drd1-cre^120Mxu^ line. This finding was surprising, as prior research by O’Neal et al. shows chemogenetic stimulation of VTA-projecting NAc neurons exacerbates heroin seeking in rats [83]. Part of this discrepancy may be due to inter-species differences in D1-MSN collaterals to the ventral pallidum (VP). In mice, over 90% of D1 MSNs that project to VTA also collateralize to VP [84]; meanwhile, O’Neal et al. report that MSNs projecting to VTA did not exhibit any collateral projections to VP in rats [83]. In VP, MSNs primarily synapse on GABAergic neurons [85], and inhibition of GABAergic VP neurons has been shown to attenuate remifentanil seeking [86]. In our study, chemogenetic stimulation of VTA-projecting MSNs is likely also affecting their collateral input to VP, thereby increasing inhibitory drive on GABAergic VP neurons that may subsequently reduce fentanyl seeking. It is likely that this effect did not express in the Drd1-cre^120Mxu^ mice because their baseline seeking responses are so low they cannot be further attenuated; additionally, it is perhaps unsurprising that chemogenetic inhibition of D1-MSNs did not restore seeking to the Drd1-cre^120Mxu^ mice considering this manipulation did not exacerbate seeking in wildtype mice. Importantly, the lack of response in the Drd1-cre^120Mxu^ mice does not reflect an inability for DREADDs to stimulate, or inhibit D1- MSNs, as we identified expected increases and decreases in *cfos* expression after DCZ treatment.

We showed that Drd1-cre^120Mxu^ mice exhibit several locomotor phenotypes, including increased locomotor response to fentanyl, reduced locomotion in a novel environment, and increased sensitivity to a D1R agonist. There are several reports on transgenic mouse lines with abnormal locomotor responses to misused drugs. For example, Drd2-EGFP mice [87] have increased striatal *Drd2* expression that reduces the locomotor response to cocaine [74]. Similarly, DAT-Ires-Cre mice (B6.SJL-Slc6a3^tm1.1(cre)Bkmn^/J, Jackson Laboratory No: 006660), which have reduced dopamine transporter (DAT) expression [88], and reduced striatal DAT function [89], exhibit attenuated amphetamine-induced locomotion [76], and similar findings were also reported in a different DAT-Cre line (Slc6a3^tm1(cre)Xz^, Jackson Laboratory No: 020080) [77]. In our study, Drd1-cre^120Mxu^ mice had elevated expression of NAc *Drd1* and a greater locomotor response to fentanyl. This effect may be driven by D1R hypersensitivity in D1-MSNs that project to VTA, where they form inhibitory synapses on GABAergic neurons [90,91] that may potentiate fentanyl-induced disinhibition of dopamine neurons. In support of increased D1R sensitivity, the Drd1-cre^120Mxu^ mice exhibited a relatively stronger locomotor stimulatory effect when given D1R agonist SKF-38393. However, the Drd1-cre^120Mxu^ mice also exhibited baseline hypolocomotion in a novel environment. Interestingly, the Drd2-EGFP line which has increased striatal D2 receptor expression [74], and the DAT-Ires-Cre line which has reduced striatal DAT function [89], both exhibit baseline hyperactivity in a novel environment [74,76]. These baseline locomotor differences are to be expected given the importance of D1- and D2-MSN signaling for normal locomotion [92].

Although the precise mechanism underlying the fentanyl-resistant phenotype of these Drd1-cre^120Mxu^ mice is unclear, looking at the transcriptional differences provides some insight. We used qRT-PCR to investigate gene expression differences in the NAc of experimentally naïve Drd1-cre^120Mxu^ mice compared to wildtype, and then compare how these genes were changed following 14 days abstinence from fentanyl self-administration. We first looked at D1-MSN markers and found Drd1-cre^120Mxu^ mice had greater *Drd1* expression than wildtypes at baseline. While wildtype mice exhibited *Drd1* downregulation following abstinence, Drd1-cre^120Mxu^ mice maintained elevated *Drd1* expression. Another D1-MSN marker, *Chrm4*, which encodes muscarinic acetylcholine receptor M4, was more highly expressed in Drd1-cre^120Mxu^ than wildtype mice at baseline, but this difference was normalized following abstinence. Muscarinic receptor activity is necessary for cholinergic modulation of heroin self-administration and seeking [93]. In the NAc, cholinergic receptors shape the expression of cue-motivated behavior via modulation of dopaminergic signaling [94]. Given that *Chrm4* expression was normalized by the 14-day seeking timepoint, it likely does not contribute to the abnormal fentanyl seeking in Drd1-cre^120Mxu^ mice. *Tac1*, another D1-MSN marker, demonstrated baseline sex differences that are consistent with findings in rat striatum [95], but no genotype differences. Following abstinence, *Tac1* was upregulated in wildtype mice, consistent with findings in morphine-exposed rats [51], and downregulated in Drd1-cre^120Mxu^ mice. *Tac1* encodes the precursor for Substance P, a neuropeptide that is particularly relevant to opioid-dependent behaviors [96], as morphine induces Substance P release in the brain [97]. Substance P is expressed in the local collaterals of NAc MSNs [98] that synapse with cholinergic interneurons containing receptors for Substance P [99,100]. This excitatory microcircuit promotes coordinated activity between D1- and D2-MSNs that shapes motivated behavior [101]. The downregulation of *Tac1* in the NAc of Drd1-cre^120Mxu^ mice likely impairs the function of this local circuit, and as such, may contribute to the blunted fentanyl seeking in these mice.

We also looked at NAc expression of D2-MSN markers, including *Drd2*, *Gpr6*, and *Adora2a*. While expression of these genes did not differ between Drd1-cre^120Mxu^ mice and wildtype mice at baseline, following abstinence Drd1-cre^120Mxu^ mice had increased *Gpr6* expression in both sexes, and increased *Adora2a* expression in males. *Gpr6,* which encodes G protein-coupled receptor 6, is enriched in NAc indirect pathway MSNs, where it plays a role in regulating instrumental conditioning [10], but a link to drug misuse has not been identified. *Adora2a* encodes adenosine A_2A_ receptors, which are colocalized on D2-MSNs [102,103], where they antagonize activity at the D2 receptor [104]. NAc A_2A_ activity has been shown to regulate cocaine seeking [105], but its role in opioid use is less studied. It is possible that these changes in D2-MSNs are compensatory, serving to oppose the changes in D1-MSNs. We also looked at expression of the enkephalin precursor gene, *Penk*, which is enriched in NAc D2-MSNs [106]. We found downregulation of *Penk*, along with what is likely a compensatory upregulation in the enkephalin receptor *Oprd1*, in post-abstinence Drd1-cre^120Mxu^ mice. Prior studies in rats suggest NAc *Penk* both influences and is influenced by opioid self-administration: *Penk* overexpression in NAc potentiates heroin self-administration [107], and NAc *Penk* expression is reduced by morphine self-administration [108]. Interestingly, our data do not follow this same pattern: Drd1-cre^120Mxu^ mice have elevated NAc *Penk* expression at baseline but do not have potentiated fentanyl self-administration, and NAc *Penk* is increased following fentanyl self-administration in wildtype mice. These disparities may be explained by differences in opioids (heroin/morphine vs fentanyl) or differences in species (rats vs mice).

In addition to the MSN markers, we also investigated other genes that, in the NAc, have known roles in opioid misuse. We found that *Ntrk2,* which encodes the BDNF receptor TrkB, was downregulated following fentanyl abstinence in wildtype mice but was unchanged in Drd1-cre^120Mxu^ mice. This finding is consistent with previous work showing morphine exposure downregulates TrkB in NAc D1-MSNs, resulting in impaired BDNF signaling that promotes morphine reward [17]. It is tempting to speculate that, given unchanged *Ntrk2* expression, Drd1-cre^120Mxu^ mice preserve normal BDNF signaling in D1-MSNs after fentanyl, thereby conferring resistance to fentanyl seeking. We also found Drd1-cre^120Mxu^ mice have blunted downregulation of *Grin2a* and *Grin2b* compared to wildtype mice following fentanyl abstinence. These genes encode the GluN2 subunits of the glutamatergic NMDA receptor. NAc glutamatergic signaling is essential for opioid seeking [109], and previous work shows chronic opioid use alters NMDA-mediated signaling in NAc [82]. The observed downregulation of NMDA receptor components in wildtype mice aligns with findings of an increased NAc AMPA/NMDA ratio that results from chronic morphine use and persists throughout protracted withdrawal [32,110]. This adaptation is believed to contribute to an increased excitatory drive onto D1-MSNs that promotes drug seeking [111]. A similar adaptation results from fentanyl abstinence [55], and this process is likely disrupted in Drd1-cre^120Mxu^ mice that do not undergo the same fentanyl-induced downregulation of NMDA subunits.

Although the precise mechanism underlying the resistance to fentanyl relapse exhibited by Drd1-cre^120Mxu^ mice is not known, it is clear that these mice have aberrant NAc D1-MSN signaling that contributes to this phenotype. Given that chemogenetic targeting of NAc D1-MSN activity did not alter fentanyl seeking in the Drd1- cre^120Mxu^ mice, it is likely that this behavior is not directly dependent on altered D1-MSN neural activity, but rather on the aberrant adaptations surrounding these neurons. The fentanyl abstinence-induced changes in NAc gene expression we observed in Drd1-cre^120Mxu^ mice suggest some of these adaptations include impaired Substance P-dependent signaling, conserved BDNF-dependent signaling, and altered NMDA plasticity, all of which could confer resistance to fentanyl seeking, although additional experiments are necessary to confirm these possibilities. A recent study proposed a model where imbalanced D1/D2-MSN plasticity promotes negative affect during opioid abstinence, and showed that restoring this plasticity through a process dependent on D1R activation attenuated negative affect and relapse [112]. Given that Drd1-cre^120Mxu^ mice exhibit increased sensitivity to D1R activation, combined with the potential adaptations to NAc plasticity discussed here, it is tempting to speculate that the Drd1-cre^120Mxu^ mice preserve D1/D2-MSN balance during fentanyl abstinence. It will be interesting for future work to investigate whether Drd1-cre^120Mxu^ mice exhibit comparable abstinence-induced negative affective behavior as wildtype mice, as the neuroadaptations described here may promote resilience through abstinence and resistance to relapse. In conclusion, our findings provide potential new molecular mechanisms and neurobiological targets for interventions to prevent relapse to fentanyl seeking.

## Acknowledgements

The authors thank Daniela Franco and Mary Kay Lobo at the University of Maryland School of Medicine for providing tissue from the FK150 Drd1-cre mouse line.

## Author Contributions

MEF designed the experiments. AM, LDF, AEO, HS, SP, and MEF performed behavioral experiments and analyzed data. AM, FAA and MEF extracted RNA and/or performed qRT-PCR experiments and analyzed data. AM and MEF wrote the manuscript and prepared the figures with input from all authors.

## Funding

This work was supported by NIH grants DA050575 and DA058661 to MEF.

## Competing Interests

The authors have nothing to disclose.

## Notes

### Competing Interest Statement

The authors have declared no competing interest.

