## Supplemental for "A Drd1-cre mouse line with nucleus accumbens gene dysregulation exhibits blunted fentanyl seeking"

### Supplemental Methods

*Drugs*

Fentanyl citrate (#22659), cocaine hydrochloride (#22165), and SKF-38393 hydrobromide (#19150), were obtained from Cayman chemical (Ann Arbor, MI). Deschloroclozapine (#7193) was obtained from Tocris Bioscience (Minneapolis, MN). All other veterinary agents (e.g. ketamine, xylazine, enrofloxacin, etc.) were obtained from PSUCOM Animal Resources. All drugs were dissolved in sterile saline (0.9%).

*Intravenous catheter surgery*

At 7-8 weeks of age, mice were implanted with indwelling jugular vein catheters (One Channel Vascular Access Button™ for Mice, Instech) under ketamine-xylazine anesthesia using previously described procedures [78]. Mice were allowed 7d to recover from surgery, during which they received daily subcutaneous injections of antibiotics (enrofloxacin 20 mg/kg) and Lactated Ringer's solution (10 mL/kg). Implanted catheters were flushed daily with 30 µL of heparinized saline containing enrofloxacin (200 IU/mL heparin, 0.227% enrofloxacin).

*Fentanyl self-administration*

Mice underwent intravenous fentanyl self-administration as previously described [78]. Briefly, mice were habituated to the operant chambers (MED Associates, Saint Albans, VT, USA) for 30 minutes one day prior to starting self-administration. Mice then underwent 10 consecutive days of fentanyl self-administration, the first 5d under a fixed-ratio 1 (FR1) schedule of reinforcement, followed by 5d under FR2 (3 h/day, 1.5 µg/kg/infusion). Sufficient responses in the active nose-poke triggered a 10 µL fentanyl infusion, turned off the house-light and the active nose-poke light, and illuminated the cue light above the active nose-poke for 1s. Any additional active responses during the 1s infusion period were recorded but did not result in further infusions ('timeout'). Responses on the inactive nose-poke were recorded but were without programmed consequences. No prior training was used for FR1 acquisition. After self-administration, all mice underwent a 1h seeking test under extinction conditions in which responses in the active nose-poke resulted in cue presentations but no drug delivery. Mice underwent the seeking test either 24h or 14d after the last self-administration session.

*Fentanyl conditioned place preference*

Conditioned place preference (CPP) was performed in a three-chambered rectangular apparatus consisting of a neutral center compartment and two side compartments with different contextual cues, connected by automatic guillotine doors, and equipped with infrared photobeam detectors that track animal position and movement (ENV-3013, MED Associates). On the first day of testing (pre-test), mice underwent a 5 min habituation in the center compartment, after which the doors opened, and they freely explored the entire apparatus for 20 min. The least-preferred chamber was subsequently assigned as the fentanyl-paired chamber. Mice then underwent 3 consecutive days of conditioning. On conditioning days, mice received an i.p. injection of saline (10 mL/kg) and were confined to the saline-paired compartment for 20 min; 4 h later, they received an i.p. injection of fentanyl (0.2 mg/kg) and were confined to the fentanyl-paired compartment for 20 min. On the fifth day of testing (post-test), after 5 min habituation in the center, they freely explored the entire apparatus for 20 min. CPP score was calculated as the difference in seconds spent on the fentanyl-paired side during post-test vs pre-test. Locomotion during fentanyl conditioning was measured as movement counts in Med-PC V Software.

*Cocaine self-administration*

A cohort of Drd1-cre$^{120Mxu}$ mice underwent intravenous cocaine self-administration as previously described [79]. Briefly, mice underwent 10 consecutive days of cocaine self-administration under FR1 (2 hours/day; 0.5 mg/kg/infusion) using a 10s timeout. Mice then underwent a seeking test 24h after the last self-administration session. This procedure was chosen to facilitate comparison between Drd1-cre$^{120Mxu}$ mice and the unstressed, pair-housed, wildtype controls from our published dataset [79].

*Locomotion*

Mice were injected with 10 mL/kg saline, or 30 mg/kg SKF-38393 s.c., then placed in the center of a Med-Associates ENV-520 open field under dim red light. Infrared beam breaks were recorded for 90 min, in 5 min block intervals, using Activity Monitor Software (Med-Associates SOF-811). Settings for distance traveled, jumps, and stereotypic movement used default software parameters.

*qRT-PCR*

Mice were euthanized by cervical dislocation, and brains were rapidly removed and chilled in ice cold PBS. Cold brains were cut into 1 mm coronal sections using an aluminum brain matrix, and tissue punches containing NAc or VTA were collected with 14-gauge needle, then snap-frozen on dry ice and stored at -80 until processing. RNA was extracted using Trizol (Invitrogen) and the RNeasy Mini Kit with a DNAse step (Qiagen). RNA concentration was measured on a Nanodrop (ND-8000, Thermo), and cDNA was synthesized with the iScript cDNA synthesis kit (Bio-Rad) using 400 ng of RNA. mRNA expression changes were measured with quantitative polymerase chain reaction (qPCR) with PerfeCTa SYBR Green FastMix (QuantaBio). Fold change mRNA was determined using the $2^{-\Delta\Delta Ct}$ method, using *Gapdh* as the housekeeping gene, and experimentally naïve, wildtype mice as the reference group (males and females combined). Primer sequences are in Supplemental Table 1.

*Chemogenetic manipulation and Stereotaxic Surgery*

Adeno-associated viruses for Cre recombinase (pENN.AAV.hSyn.HI.eGFP-Cre.WPRE.SV40), Cre-dependent Gq-coupled (AAV9-hSyn-DIO-hM3D(Gq)-mCherry) or Gi-coupled (AAV9-hSyn-DIO-hM4D(Gi)-mCherry) Designer Receptors Exclusively Activated by Designer Drugs (DREADDs), and mCherry control (AAV9-hSyn-DIO-mCherry) were acquired from Addgene (viral preps # 105540-AAVrg, 44361-AAV9, 44362-AAV9, 50459-AAV9).

Mice were anesthetized with 1-4% isoflurane gas in oxygen delivered at 1 liter/min, and affixed in a stereotaxic frame (Stoelting, Dale IL). A small burr hole was made over the NAc and/or VTA, using the following coordinates (in mm, relative to bregma): NAc (AP +1.6, ML +1.5, DV -4.4), VTA (AP -3.2, ML +1.0, DV -4.6). Viral vectors were delivered with Hamilton neurosyringes with 33-gauge needles at a rate of 100 nL/min for a total volume of 300 nL/virus, then needles were left in place for 5 min to minimize spread up the tract. For Drd1-cre$^{120Mxu}$ mice, Cre-dependent DREADDs were infused bilaterally in NAc. For wildtype mice, retrograde Cre was infused bilaterally in VTA, and Cre-dependent DREADDs infused bilaterally in NAc.

For chemogenetic manipulations during fentanyl seeking, stereotaxic surgery was performed 24 h after the last self-administration session, allowing 14d for viral expression before the seeking test. DREADDs were activated via i.p. injection of deschloroclozapine (DCZ, 0.1 mg/kg) 20 min before the seeking test. Mice were perfused after seeking and brains were processed for immunohistological verification of DREADDs expression in NAc. For chemogenetic manipulation during fentanyl CPP, stereotaxic surgery was performed on naïve Drd1-cre$^{120Mxu}$ mice, and they began CPP 14d later. Mice were injected with saline (or DCZ) 20 min before the post-test, then 4h later, injected with DCZ (or saline) 20 min before a second post-test. To capture c-Fos mRNA expression resulting from DREADDs activation, a subset of mice were euthanized via cervical dislocation immediately following the second post-test while DCZ was still on board. NAc and VTA punches were collected as above from mice with bilateral mCherry expression, visualized with a stereomicroscope equipped with a fluorescence adapter (NightSea, Hatfield PA).

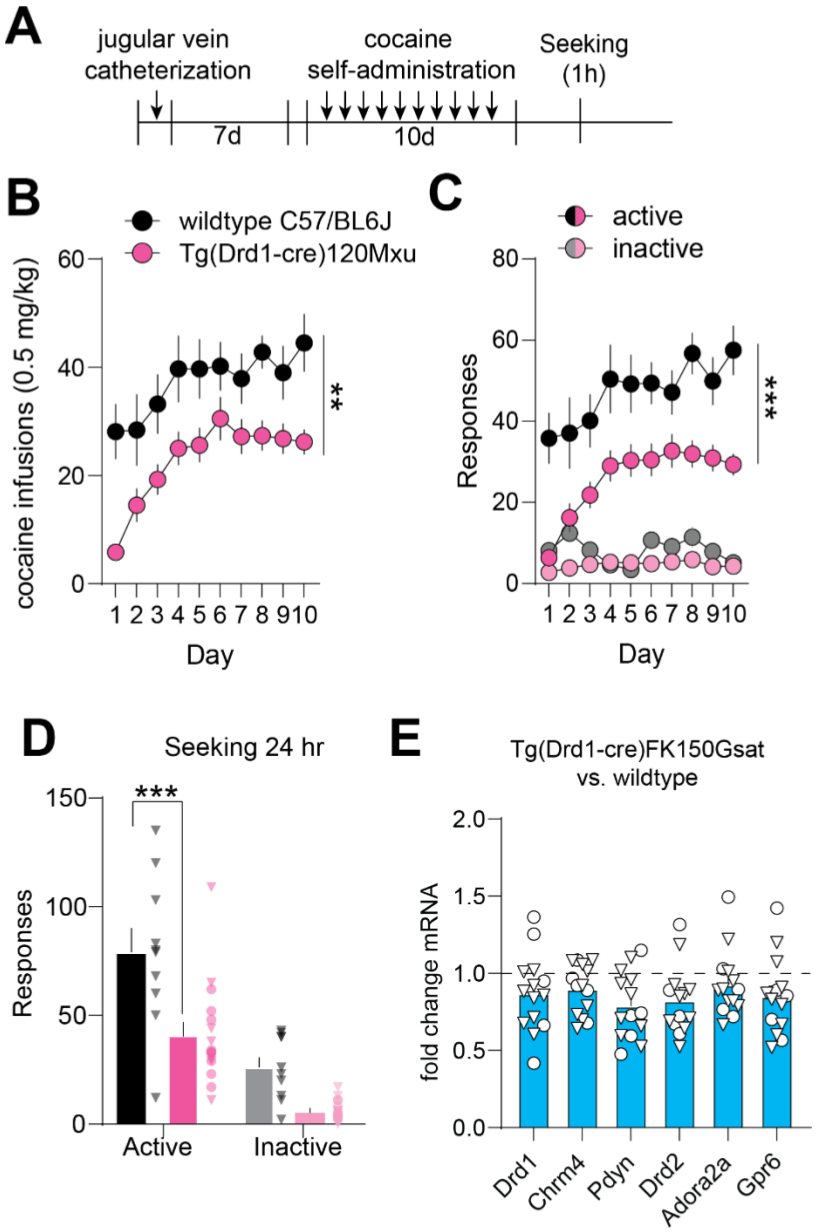


**Supplemental Figure 1.** (**A**) Experimental timeline for cocaine self-administration experiments. Following recovery from jugular catheterization surgery, mice underwent 10 days of cocaine self-administration training (0.5 mg/kg/inf) under FR1 as in our published work [79] **(B**) Number of cocaine infusions earned during self-administration training in wildtype (black) and Drd1-cre$^{120Mxu}$ (magenta) mice. **, p=0.0031, main effect of genotype in RM-ANOVA. (wildtype n=10♂; Drd1-cre$^{120Mxu}$ n=9♀, 7♂) (**C**) Number of active and inactive responses during cocaine self-administration training in wildtype and Drd1-cre$^{120Mxu}$ mice. ***, p=0.0003, main effect of genotype. (**D**) Number of active and inactive responses during a non-reinforced seeking task 24 hr after the last self-administration session. ***, p=0.0001, Sidak's post-hoc after 2-way ANOVA. (**E**) Expression of D1- and D2-MSN markers in NAc of a different Drd1-cre line (Gensat FK150, n=5♀, 9♂). Hashed line is average expression wildtype mice (male and female combined). Data are presented as mean±SEM.

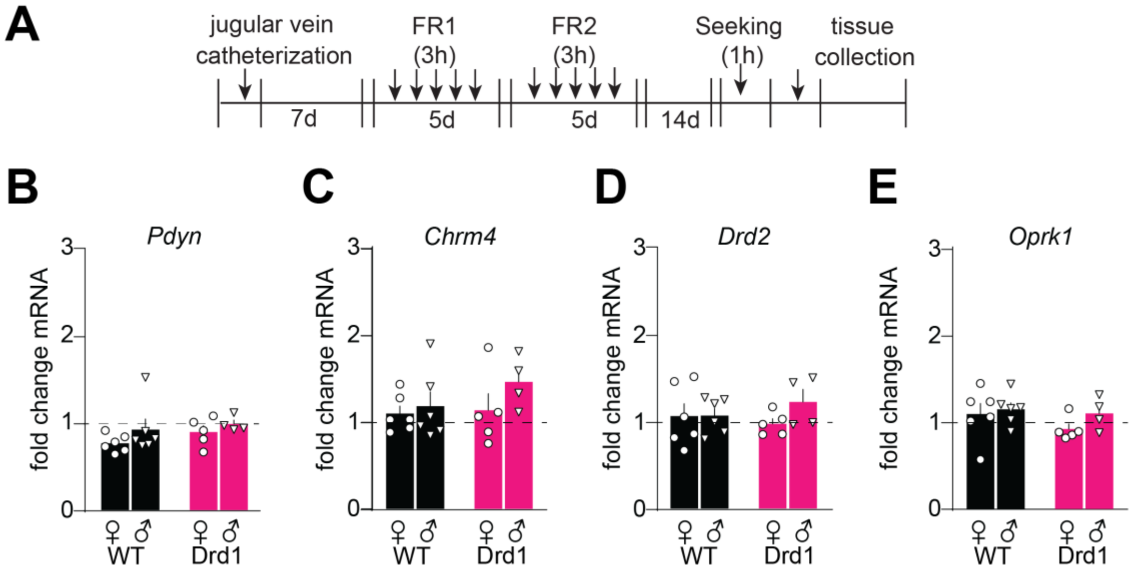


**Supplemental Figure 2.** Fentanyl self-administration normalizes some gene expression differences between Drd1-cre$^{120Mxu}$ and wildtype mice. (**A**) Experimental timeline. (**B-E**) Fold change mRNA relative to experimentally naïve male and female wildtype mice (dashed line; wildtype n=6♀, 6♂; Drd1-cre$^{120Mxu}$ n=5♀, 4♂). Data are presented as mean±SEM. (**B**) Preprodynorphin. (**C**) Muscarinic Acetylcholine receptor M4. (**D**) Dopamine receptor D2. (**E**) Kappa opioid receptor.

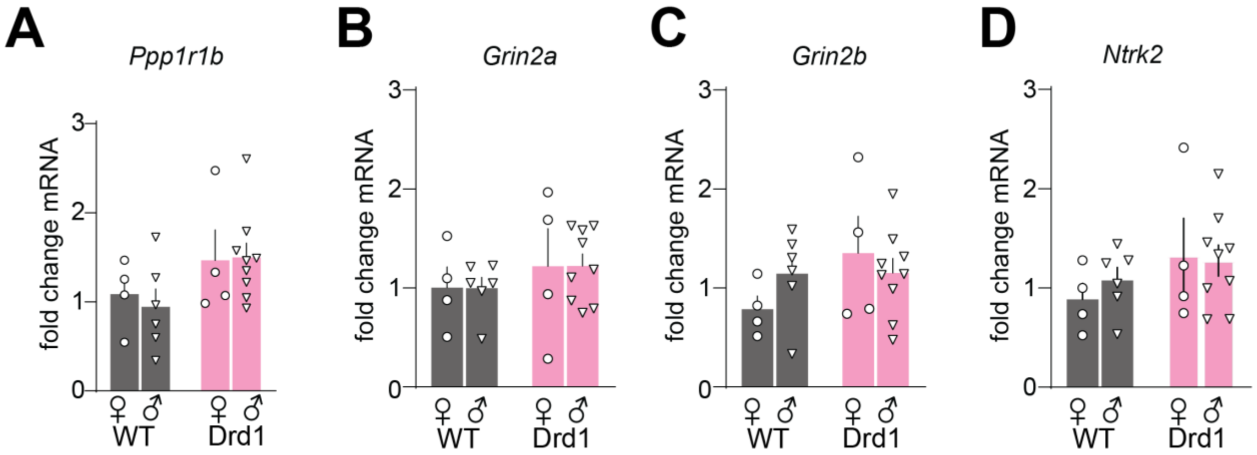


**Supplemental Figure 3.** Fentanyl-influenced genes are unaltered at baseline in Drd1-cre$^{120Mxu}$ vs wildtype mice. Fold change mRNA in experimentally naïve wildtype (black) and Drd1-cre120Mxu (magenta) mice, relative to average of naïve male and female wildtype mice (wildtype n=4♀, 6♂; Drd1-cre120Mxu n=4♀, 9♂). Data are presented as mean±SEM. (**A**) DARPP-32. (**B**) NMDAR subunit 2a. (**C**) NMDAR subunit 2b. (**D**) TrkB receptor.

| Gene | Forward Primer | Reverse Primer |
|---|---|---|
| Drd1 | GAGCGTGGTCTCCCAGAT | GGATGCTGCCTCTTCTTCTG |
| Chrm4 | ATCGGCTACTGGCTCTGCTA | TACTGGCACAGCAAAAGGTG |
| Pdyn | CTCCTCGTGATGCCCTCTAAT | AGGGAGCAAATCAGGGGGT |
| Tac1 | TGTTGGACTAATGGGCAAAA | GATAGTGCGTTCAGGGGTTT |
| Drd2 | TCAGATGCTTGCCATTGTTC | GTGAAGGCGCTGTAGAGGAC |
| Adora2a | CACGCAGAGTTCCATCTTCA | AATGACAGCACCCAGCAAAT |
| Gpr6 | GAGGATAGCCAGGCACACAG | ACCACTTGGGACTCGTTGAG |
| Penk | GAGAGCACCAACAATGACGAA | TCTTCTGGTAGTCCATCCACC |
| Oprm1 | CCAGGGAACATCAGCGACTG | GTTGCCATCAACGTGGGAC |
| Oprd1 | CCATCACCGCGCTCTACTC | GTACTTGGCGCTCTGGAAGG |
| Oprk1 | TCCCCAACTGGGCAGAATC | GACAGCGGTGATGATAACAGG |
| Oprl1 | CGTGCCCTTGATGTTCGGA | GGCCCCAATAGTCCTGAGG |
| Ppp1r1b | CCAACCCCTGCCATGCTTT | TTGGGTCTCTTCGACTTTGGG |
| Ntrk2 | TTGTGTGGCAGAAAACCTTG | ACAGTGAATGGAATGCACCA |
| Grin2a | ACGTGACAGAACGCGAACTT | TCAGTGCGGTTCATCAATAAC |
| Grin2b | GCCATGAACGAGACTGACCC | GCTTCCTGGTCCGTGTCATC |
| Gapdh | AGGTCGGTGTGAACGGATTTG | TGTAGACCATGTAGTTGAGGTCA |

**Supplemental Table 1.** Primer Sequences for qRT-PCR.
